# Responses of tree species traits to soil variation in the cerrado

**DOI:** 10.1101/2020.02.20.955955

**Authors:** João Augusto Alves Meira-Neto, Glaucia Soares Tolentino, Tillmann Buttschardt, Cristina Máguas

**Affiliations:** Laboratory of Ecology and Evolution of Plants – LEEP/PPGBot-UFV, Universidade Federal de Viçosa, Campus UFV s/n, Viçosa, State of Minas Gerais, Brazil, 36570-000; Centre for Ecology, Evolution and Environmental Changes – CE3C, Faculdade de Ciências da Universidade de Lisboa, Edifício C2, Lisboa, Portugal, 1749-016; Institute of Landscape Ecology - ILÖK, University of Münster, Münster, Germany, D-48149

**Keywords:** Nitrogen-fixing, Cerrado, functional traits, aluminium saturation, water use efficiency

## Abstract

**Aims:** The Cerrado is a rich tropical savanna in plant species and to understand how key functional traits respond to soil variables can help to understand this hotspot functioning. This work aimed to answer i) whether the Cerrado tree species respond to soil by functional traits, ii) how the functional traits respond to soil variation in the Cerrado, and iii) whether the functional traits responses are interconnected.

**Methods:** We used an RLQ method to associate soil variables to functional traits and GLMs for additional analysis. We used the nitrogen-fixing character as categorical trait and maximum plant height, maximum stem diameter, stem elongation, leaf nitrogen, leaf carbon, leaf C/N, leaf d15N and leaf d 13C as continuous traits.

**Results:** The RLQ showed that tree species responded to the soil variables with differences between nitrogen-fixing species and other species. The increase of CEC and decrease of aluminium saturation induced an increase of SLA and of stem elongation. CEC and aluminium saturation did not influence the leaf N% and C%. The increase of soil inorganic nitrogen is associated to an increase of leaf N% and of leaf C%. The C/N ratio explained negatively the δ13C and the stem elongation variation. Nitrogen fixing species presented low C/N ratios and high δ13C.

**Conclusions:** The relative disassociation of the variation of SLA and stem elongation (mostly driven by CEC and aluminium variation) from the variation of N% and C% (mostly associated with inorganic nitrogen variation) can be a result of enhanced water use efficiency in N-enriched plants.

## Introduction

The Cerrado is the largest savanna in the Neotropics, a biodiversity hotspot for plants (Simon et al. 2009; Souza□Neto et al. 2016; Françoso et al. 2019) and strongly influenced by many soil variables (Rossatto et al. 2012; Franco et al. 2014). Despite some studies have demonstrated the soil-plant relationship in the Cerrado (de Assis et al. 2011; Neri et al. 2012; Meira-Neto et al. 2017). the studies that evaluate the specific response of plant functional traits to the soil properties variability are still limited. In such complex hotspot, to decrease the information from the high number of variables and species to few key functional traits responding to variables can help to understand the environmental influence on plants (Dıa z and Cabido, 2001; Lavorel and Garnier, 2002).

Functional traits are key characteristics that allow trees to deal with the environment (Baraloto et al. 2012). The specific leaf area (SLA) is a synthetic leaf trait that responds congruently throughout different biomes, climates and latitudes to resource availability (Reich et al. 1997). SLA is positively related to light use efficiency, to productivity, to competitive ability and negatively related to stress tolerance and to leaf lifespan (Wright et al. 2001; Valladares and Niinemets 2008). Stem elongation expressed as a ratio of maximum plant height to maximum stem diameter (Hmax/Dmax) is a trait positively related to light competition ability (Bagousse□Pinguet et al. 2017; Candido et al. 2019) and to resource availability (Goldstein et al. 2019; Khan et al. 2020). The leaf contents of nitrogen (N%), carbon (C%) as well as the carbon to nitrogen ratio (C/N) indicate the nutrient status for plants and the stress level. The N% and C% are positively related to nutrient status of leaves and the C/N is positively related to the nutrient limitation (Cornelissen et al. 2003; Pérez-Harguindeguy et al. 2013). Together, the traits separately or combined as ratios should show the main strategies of plants responding to soil variability in the Cerrado with leaf (SLA), stem (Hmax / Dmax) and root (nutritional status) functioning in a comprehensive analysis (Westoby 1998) of the environment in which the Cerrado plants are found.

Besides the continuous functional traits, functional categories have been used for the tree species of Cerrado. The nitrogen-fixing leguminous can be understood as a functional group of trees species that occur throughout Cerrado physiognomies and dominate the dense woodlands on acidic dystrophic soils called Dystrophic Cerradoes (Meira-Neto *et al*., 2017), but little is known besides the nitrogen-fixing trait. Dystrophic Cerradoes are dominated by nitrogen-fixing leguminous trees (Goodland 1971; Oliveira-Filho and Ratter 2002; Meira-Neto et al. 2017) and have high abundance of trees, with much higher biomass than open Cerrados, occur on deep acidic dystrophic soils with high levels of Al^3+^ and with high levels of inorganic nitrogen.

Functional ecology in the Cerrado has described vegetation functioning (e.g., Franco et al. 2014; Meira-Neto et al. 2017) and ecological processes (Freitas et al. 2012) by models. Thus, models are tools for the Cerrado understanding in different scenarios, but the soil complexity in a plant diversity hotspot hinder the functional understanding blurring the plant-soil relations. The Cerrado complexity can be disentangled by the use of by functional traits of plants. Few vegetative traits of leaf (e.g., specific leaf area - SLA) and stem (e.g., stem elongation) as well as the percentage of carbon and nitrogen and their stable isotopes in plants are key traits (Westoby 1998; Máguas and Griffiths 2003; Franco et al. 2014; Meira-Neto et al. 2017, 2018) related to plant growth and can help to understand the general functional effects of soil on plants.

This research aimed to answer i) whether the tree species in the Cerrado respond to soil variation by responsive functional traits, ii) how the studied functional traits respond to soil variation in the Cerrado, and iii) if the functional traits responses to soil variation are interconnected or disconnected from each other in the Cerrado. We studied a Cerrado with a wide range of soil characteristics within an area without climatic, topographical and dispersion limitation biases and with negligible disturbances, a suitable system for increasing knowledge regarding plant functional responses to soil variation.

## Material and methods

### Study area

The study was carried out in the Paraopeba Reserve with an area of 200-ha in the state of Minas Gerais (19°20’S, 44°20’W), Brazil. The climate of the region is classified as Aw (tropical humid) by the Köppen system with a rainy summer from October to April and a dry season from May to September. The mean annual temperature and mean annual rainfall are 20.9°C and 1328 mm, respectively. The vegetation of the reserve is the product of regeneration after clear-cutting in 1952. There are records of fires in 1960 and 1963, since which the area as a whole has been protected from fire (Neri et al. 2012).

We carried out the study on a wide range of soils spread over a short distance, where differences in vegetation are ascribed to soil attributes since variation in climate and topography was negligible (Neri et al. 2012).

The soils are classified as i) dystrophic Haplic Cambisol, ii) Yellow Latosol, iii) Red-Yellow Latosol, iv) dystrophic Red Latosol and v) mesotrophic Red Latosol (EMBRAPA 2006; Neri et al. 2012). Along this gradient the dark-redder the soil the greater the biomass and density of woody plants. Colluvial materials from weathered limestone or *in situ* alteration of such calcareous rocks influence both eutrophic and dystrophic red soils. On the other hand, Cerrado savannas are related to either shallow or deep yellowish soils (Cambisols/Latosols) developed from slate, a pellitic Al-rich nutrient-poor metamorphic rock. Since there is a wide range of soil characteristics within a small area, this fragment of Cerrado is a suitable system for increasing knowledge regarding plant functional groups with different strategies for resource use and stress tolerance without climatic, topographical and dispersion limitation biases.

### Sampling plants and soil

We used 75 plots (20-m x 20-m) in the Paraopeba Reserve in order to know which woody species were the dominant. As 75% of dominance is the lower limit for the highest level of dominance for sampling vegetation methods as Braun-Blanquet and Domin-Krajina (Mueller-Dombois and Ellenberg 2003; Maarel 2009), we considered as rulers of the Cerrado functioning the species that collectively comprise ∼ 75% (i.e., 74.06%) of the abundance. We used data from a previous survey on 3 hectares with 15 plots of 20-m x 20-m on each soil type, 75 plots in total. We found 14,671 individuals of 174 woody species of 51 families in the woody communities of the studied vegetation. Of these, 10,866 individuals belonged to the 34 species that made up 74.06% of the total abundance of the community. We used the 75 plots only to choose that 34 species with ∼75% of abundance. From those 75 plots, we only used 24 plots to analyze soils, light and to take leaf samples of the most abundant species (i.e., dominant species).

The structure of the woody plants was evaluated by equally distributing 24 plots (20-m x 20-m) along the aforementioned soil gradient. We sampled three plots of Inceptisol (Cambisol), three plots of mesotrophic Red Latossol, three plots of dystrophic Red Latossol, six plots of Yellow Latosol and nine plots for Red-Yellow Latossol (Yellow Latosol and Red-Yellow Latosol are intermediary soils with more plots to better represent the intermediate portion of the gradients). All plots were randomly selected inside soil types areas. The soil characterization is in previous publications (Neri et al. 2012, 2013). From July 2011 to September 2012 we recorded all individual woody plants with stem circumferences equal to or greater than 10 cm at ground level as usual in Cerrado. The classification of species into families followed APG IV (The Angiosperm Phylogeny Group 2016), with the nomenclature of species and abbreviations being in agreement with Brazilian Flora Checklist (www.floradobrasil.jbr.gov.br).

Soil samples were collected from all 20-m x 20-m plots, each one comprising 10 merged subsamples from a depth of 0–10 cm. Soil samples taken for chemical analysis were air-dried and sieved. Since Cerrado soils generally possess low nutrient status and fertility, which have been shown to influence its vegetation, we used cation exchange capacity (CEC), inorganic N and pH as proxies for soil fertility. Furthermore, since aluminium saturation determines the fraction of Al^3+^ in CEC, it was considered a main soil factor. The pH values were obtained from aqueous solution while inorganic N was gathered from the sum of nitrate (NO^3-^) and ammonium (NH_3_) content. Colorimetric analyses were used for measuring NO^3-^ and NH_3_. All analyses were performed in the Soil Laboratory at the Universidade Federal de Viçosa, following methods for Brazilian tropical soils (EMBRAPA, 1997).

### Functional traits

We evaluated functional traits of 34 most abundant species, which accounted for 74.06% of the cumulative relative abundance in the Cerrado of Paraopeba Reserve. All individuals of selected species were assessed and sampled for measuring traits. The functional traits were calculated from direct measurements from each one of the 3796 sampled individuals.

Trait-based analysis requires selection of critical traits to the community processes of interest (Kraft et al. 2007). Concerning the Cerrado, key functional traits should enable plants to grow in stressing conditions in poorer soils without light competition and to compete in less stressing soils, but with more light competition. Six key traits (Table 1) related to competitive ability, resources use and tolerance were used (Cianciaruso et al. 2012; Pérez-Harguindeguy et al. 2013). The traits were measured by widely accepted methods (Cornelissen et al. 2003; Pérez-Harguindeguy et al. 2013). The sampling was plot-specific, which allowed us to assess environmental factors (Cianciaruso et al. 2009). We used the tag nitrogen-fixing for nodulating leguminous trees (Sprent 2009; Meira-Neto et al. 2017) for sampled species (Appendix S1).

**Table 1.**
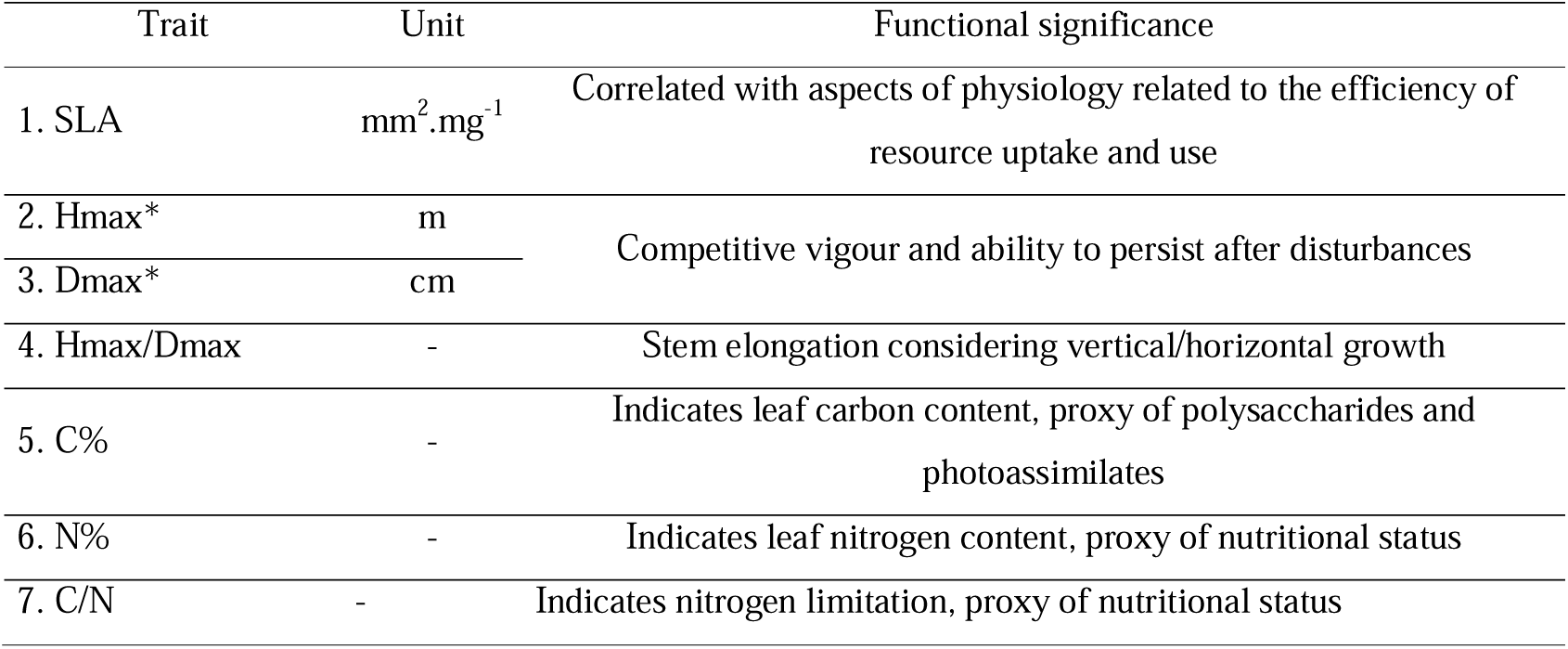
Plant traits used to analyze functional groups of a Cerrado community at Paraopeba Reserve, Minas Gerais. (*) indicates a trait that was not directly used in the data analysis, but which is part of a rate. SLA = specific leaf area, Hmax = maximum height, Dmax = maximum diameter, C/N = carbon to nitrogen ratio of leaf.

### Carbon, nitrogen and stable isotopes analyses (δ^13^C and δ^15^N)

We used stable isotopes analyses in order to understand the conditions of functional groups and set of species in each plot under different soil and light conditions. Three leaves healthy and fully developed from all individuals sampled in each plot were taken during the rainy season (March of 2013) totalizing 3796 individuals. The leaves were sampled at a similar distance from the ground and at the north side of the canopy. The leaves were dried to constant weight (65°C) without petioles and midribs and ground to a powder using a ball mill (Retsch, Haan, Germany) for measuring carbon, nitrogen, δ^13^C and δ^15^N. Nitrogen and carbon concentration and δ^13^C and δ^15^N were analysed using an elemental analyzer (HEKAtech, Weinberg, Germany) with a continuous-flow stable isotope ratio mass spectrometer (ISOPRIME, GV, Manchester, UK)and measured against an ammonium sulphate standard (IAEA.N2). C and N isotope ratios were presented in δ notation:

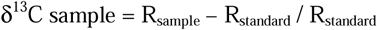

where R_standard_ is the ^13^C/^12^C of Pee Dee Belemnite (PDB) and R_sample_ is the ^13^C/^12^C ratio of the sampled leaves.

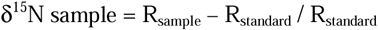

where R_standard_ is the ^15^N/^14^N ratio of atmospheric N_2_ and R_sample_ is the ^15^N/^14^N ratio of the sampled leaves. The repeated measurement precision was 0.2 ^0^/_00_.

### Responses of functional traits

In order to encompass species that are placed similarly along the gradient and that were functionally tagged, we constructed a numerical matrix, with averaged values of SLA, ratio of maximum height to maximum diameter (Hmax/Dmax) as stem elongation measurement, leaf N% and leaf C%, as individual-weighted means in plots for the 34 dominant species (Appendix S1). We did not use the C/N because of the redundancy with leaf N% and leaf C%. We used the tags of nitrogen-fixing and other for the sampled species. In order to correlate to test directly the soil influences on the functional responses we performed a predictive fourth-corner approach proposed by (Brown et al. 2014) combined with RLQ analysis that integrate correspondence analysis, Hill ordination ad PCA to produce a canonical ordination with the dispersions of species, of environmental variables and of functional traits respectively (Dray et al. 2014). RLQ analysis was carried out using the functions ‘dudi.coa’, ‘dudi.hillsmith’, ‘dudi.pca’ and ‘rlq’ of the ade4 package (Dray et al. 2018) for canonical ordination of species abundances, functional traits (SLA, Hmax/Dmax, %C and %N), plots and three soil variables (CEC, aluminium saturation and inorganic nitrogen, all square-rooted transformed). The Fourth-corner analysis was carried out using the ‘fourthcorner’ function of the Ade4 package performing 50 000 permutations with FDR adjustment of P values in order to avoid type I error for multiple comparisons between the soil variables and the functional traits of the RLQ analysis (Peres-Neto et al. 2017; Braak et al. 2018). All analyses were performed within the R statistical environment (R Development Core Team, 2015).

In order to show how the functional traits are associated to each other, we performed global GLMs with each trait as response variable against the others as explaining variables (leaf N %, leaf C %, leaf C/N, SLA, stem elongation, δ13C and δ15N) using the ‘glm’ function of the lme4 package with gaussian error distribution (R Development Core Team 2015). We used the ‘dredge’ function from the MuMIn package (Bartón 2018) to select the combinations of three or less uncorrelated predictor variables with r < 0.6 according to a protocol (https://github.com/rojaff/dredge_mc) to avoid overfitting since we have measurements for 34 species. Best models were selected using AIC (Symonds and Moussalli 2011), and all selected models with ΔAIC lower than 2 were considered equally good. When more than one model was selected, we calculated a conditional model-averaged estimates using the ‘model.avg’ function from the MuMIn package with the significance of each predictor evaluated using likelihood ratio tests (Bartón 2018).

## Results

The canonical RLQ ordination showed that tree species responded to the soil variables. Nitrogen-fixing species were associated with higher carbon and nitrogen percentages in their leaves while other species were mainly ordinated along the axis associated to stem elongation and to SLA variation with *Magonia pubescens* at the score of the highest values of SLA and stem elongation and *Salvertia convallariodora* at the score of the lowest values of SLA and stem elongation (Figure 1).

**Figure 1.**
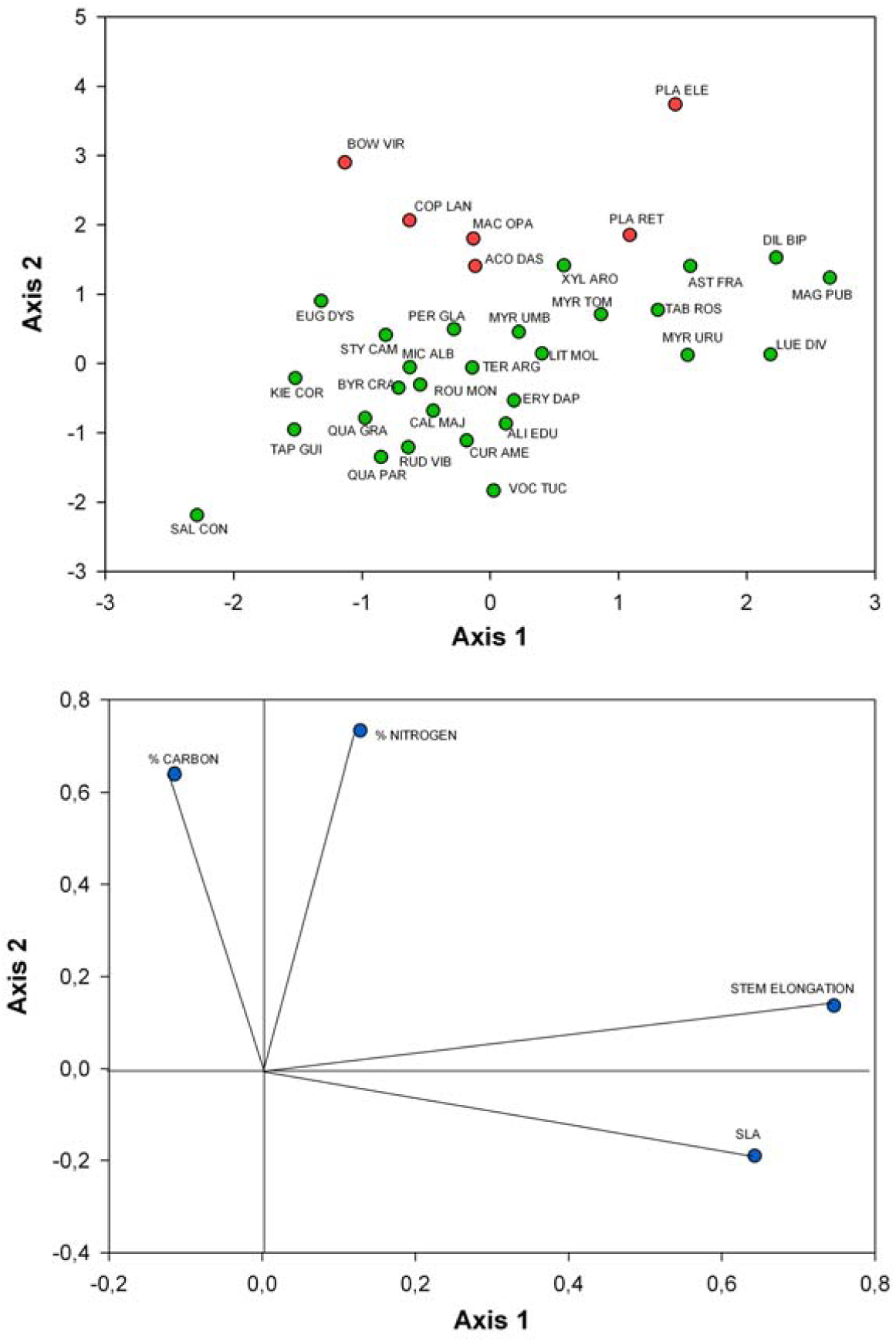
Canonical RLQ ordination of the dominant 34 woody species and environmental variables CEC, aluminium saturation and soil inorganic N; red dots are nitrogen-fixing species, green dots are other species. SLA – specific leaf area, elongation – maximum height to maximum diameter ratio, % nitrogen – leaf nitrogen content (%), % carbon – leaf carbon content (%). Species codes in Appendix S1 and Appendix S2.

The fourth-corner analysis showed that the CEC in soil is positively related to SLA and to stem elongation (Hmax/Dmax) on Cerrado trees. On the other hand, the aluminium saturation in soil is negatively related to SLA and to stem elongation on Cerrado trees (Figure 2) unveiling an opposite effect of CEC. CEC and aluminium saturation did not influence the leaf N% and C%. The soil inorganic nitrogen is positively related to leaf N% and to leaf C% (Figure 2) according to the fourth-corner analysis.

**Figure 2.**
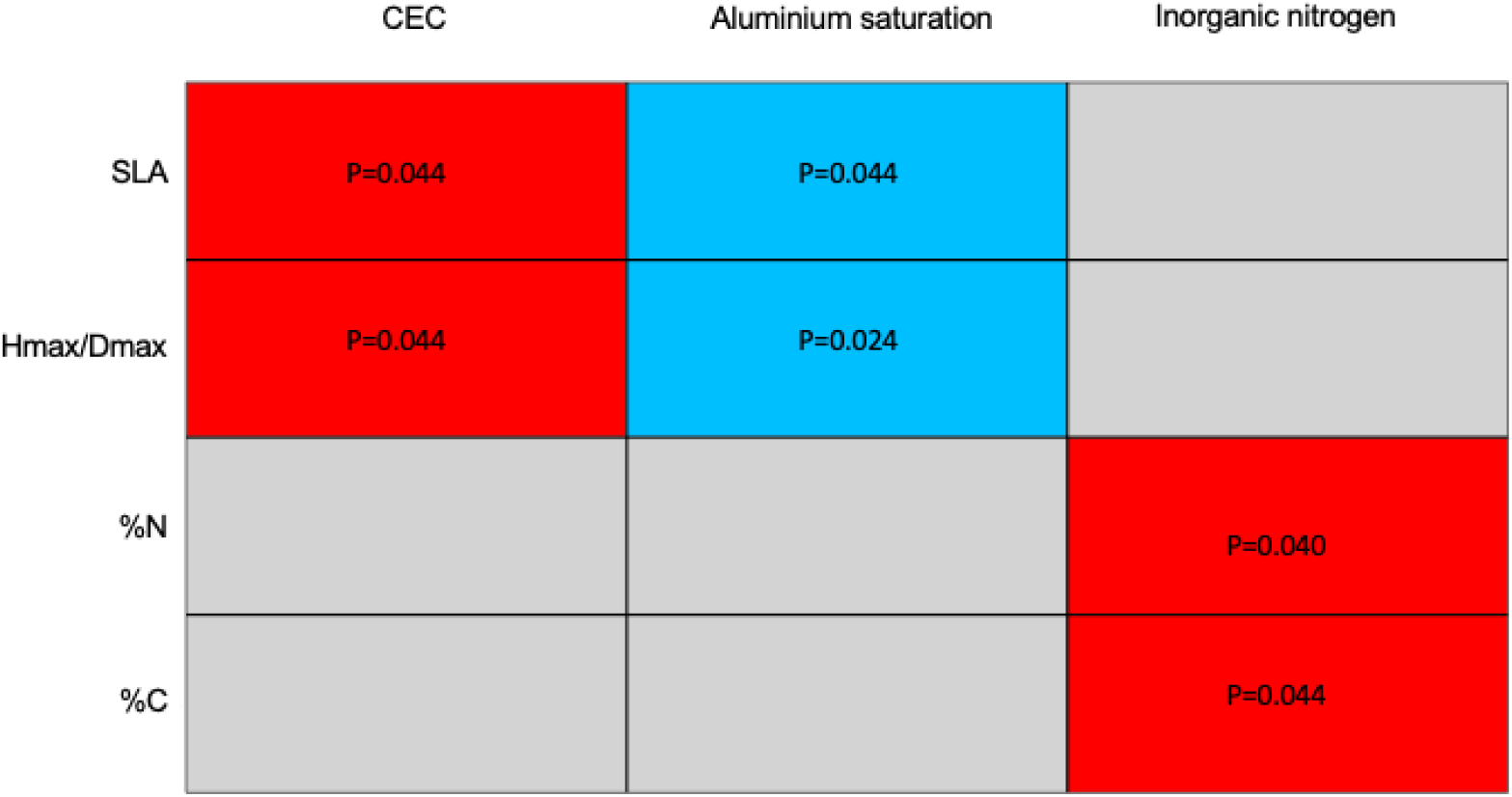
Results of the fourth-corner analysis. In red, P-values of positive relations between Cation Exchange Capacity (CEC) and specific leaf area (SLA); between CEC and stem elongation (Hmax/Dmax). In blue, P-values of negative relations between aluminium saturation (m) and specific leaf area (SLA); between aluminium saturation and Hmax/Dmax (stem elongation).

The GLMs shown that stem elongation responded also positively to leaf N% and negatively to leaf C/N showing that stem elongation is somehow associated with leaf nitrogen status (Figure 3A). SLA, d13C, leaf C% and d15N were also associated to stem elongation variation but were not significant in the averaged models. The leaf C/N is negatively related to δ13C (Figures 3 and 4) despite δ15N, stem elongation, C% and SLA also appear in the selected models without significance in averaged models (Figure 3B). The leaf C% as well as C/N were associated to leaf N% (Figure 3C). Leaf N% and C% obviously explained C/N (Figure 3D). The nitrogen fixing species presented low C/N ratios and high δ13C proportions (Figure 4).

**Figure 3.**
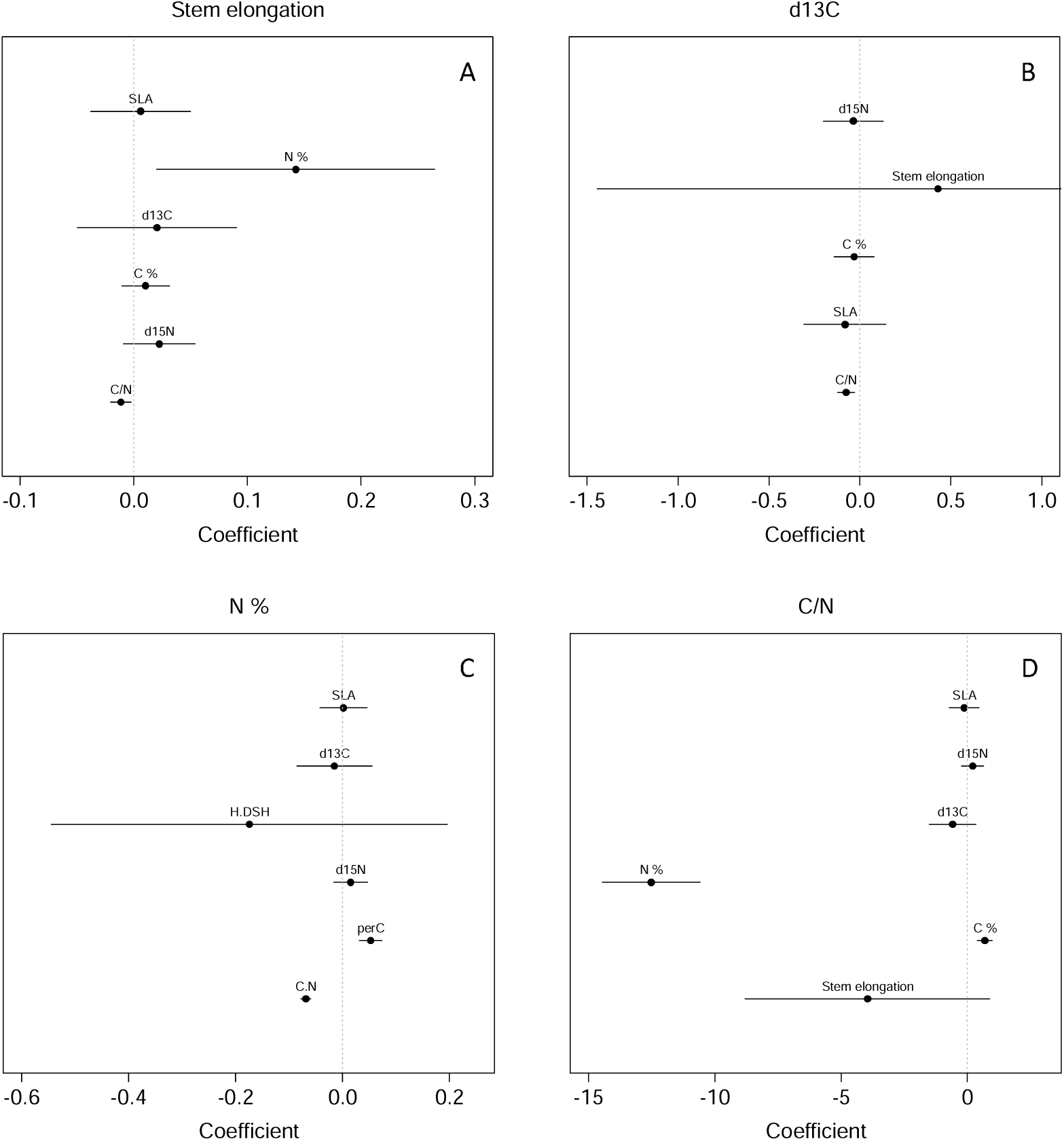
Responses of functional traits of tree species to soil variation in the Cerrado. A - stem elongation responses, B-δ13C (d13C) responses, C - leaf nitrogen (N%) responses and D - C/N responses. Confidence interval = 95%.

**Figure 4.**
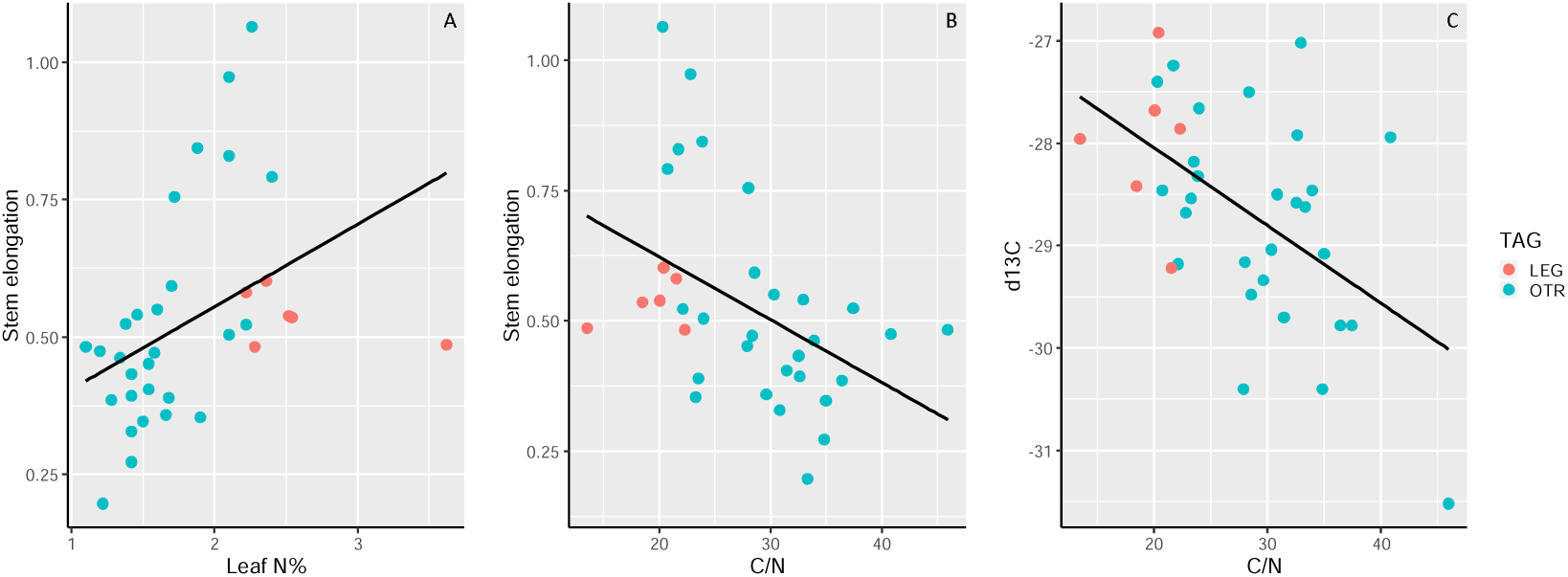
Regressions of two global GLMs, the first of leaf N % and leaf C/N predicting stem elongation (A and B, p=0.0222 and p=0.0137 respectively) and the second of C/N predicting leaf δ^13^C (d13C) (C, p = 0.00134) of the 34 Cerrado tree species. Species tags: nitrogen-fixing leguminous species (LEG), other species (OTR). LEG has lower C/N (p=0.00054) and higher Leaf N% (p= 4.66e-06) than OTR tested by single GLMs.

## Discussion

SLA and stem elongation were the functional traits that responded to the variation of CEC and of aluminium saturation in soil. The more fertile soils with higher CEC and lower aluminium saturation cause elongated stems as well as cause high SLA expected in more productive ecosystems (Reich et al. 1997; Westoby 1998). Stem elongation also positively associated to leaf nitrogen status. Leaf N% content and leaf C% were positively associated with soil inorganic nitrogen, all of them possibly interconnected in a two-way nitrogen exchange between the soil pool and the biomass pool enhancing the photosynthetic activity that captures atmospheric carbon (Wright et al. 2003; Bustamante et al. 2006). The CEC/aluminium saturation influences SLA/stem elongation rather independently from the influence of soil inorganic nitrogen on leaf N% and leaf C%.

The high SLA (i.e., short leaf lifespan) is associated with some species such as *Magonia pubescens, Luehea divaricata* and *Dilodendrum bipinatum* that are commonly found in tropical dry forests (Oliveira-Filho et al. 2006) and suggests high productivity (Reich *et al*., 1997) together with high stem elongation. Nitrogen fixers presented high leaf N%, high C% and intermediate situations of SLA and of stem elongation in the studied Cerrado. The moste elongated species dominate the most fertile soils with high CEC (i.e., Mesotrophic Red Latosol) and the nitrogen fixers dominate the poorer Dystrophic Red Latosol with medium-to-high CEC (Meira-Neto et al. 2017). The SLA and stem elongation are functional traits that are related to competitive ability for light, to long lifespan and to habitats with high productivity (i.e., high SLA; Westoby 1998; Moles et al. 2009; Pérez-Harguindeguy et al. 2013; Kunstler et al. 2016). Therefore, species with higher SLA and more elongated stems seem to be more competitive than nitrogen-fixers in soils with higher CEC and lower aluminium saturation. On the other hand, nitrogen-fixers dominate Dystrophic Cerradoes possibly using their N-enriched leaves to be more efficient in the CO_2_ use (i.e., more C%) in that very acidic soil.

Our RLQ results reinforce the CEC as a synthesis of soil fertility explaining plant responses in Cerrado (Neri *et al*., 2013) and reinforce the understanding that exchangeable aluminium is a main source of stress for woody plants in tropical savannas on acidic soils (Furley and Ratter 1988; Motta et al. 2002; Sugihara et al. 2014). The exchangeable aluminium is not only a stressor in the Cerrado but also in many other vegetations worldwide and the results found for Cerrado can be found in other vegetations on acidic soils (see Watanabe and Osaki 2002; Kochian et al. 2004). According to our findings, soils with low CEC and with toxic levels of Al^3+^ are associated with less elongated stemmed trees with lower SLA and additionally can cause open physiognomies (see Goodland and Pollard, 1973; Ruggiero *et al*., 2002; Neri *et al*., 2012) with less biomass and less abundances of woody plants (Silva *et al*., 2013) worldwide.

Nitrogen-fixing trees were associated to high leaf contents of nitrogen and carbon. Their low leaf C/N and high leaf N% are congruent with the nitrogen-fixing function (Bustamante et al. 2004, 2006) as well as with the high inorganic nitrogen contents in soil. This relation is possibly interconnected in a two-way nitrogen exchange between the soil nitrogen pool and the biomass nitrogen pool. From the leguminous species of this study, only *Copaiferea langsdorffii* may not be a nitrogen-fixer (Sprent 2009) but behave as the other leguminous tree species in our analyses and deserves attention in future studies on nitrogen-fixing species.

Despite the reported evidence of nitrogen-fixing by nodulated genera in the Cerrado (Sprent *et al*. 1996), our results of leaf δ^15^N were not significantly related to the studied functional traits. The proportion of this stable isotope should be further studied in soil and plants to determine whether the sources of N for the different species are the same (Marshall et al. 2007; Craine et al. 2015) and whether there are ecological processes involved in nitrogen uptake by non-nitrogen-fixing plants.

The negative relation found between C/N and δ^13^C show that the lower the N nutritional status, the lower the ^13^C proportion in leaves. A possible explanation of very low δ^13^C in some species may be that deep roots can access water during the dry season on deep soils in open Cerrado (Rawitscher et al. 1943; Ferri 1944; Rachid 1947; Rossatto et al. 2012). Therefore, low ^13^C proportion might be caused by high stomatal conductance because high CO_2_ assimilation lowers the δ^13^C in biomass as Rubisco freely discriminates and prefers ^12^C to ^13^C (Kohn, 2010). High N% is associated with high δ^13^C because higher nitrogen status in leaves may enhance drought tolerance (Wright et al. 2001, 2003) and should be further investigated in nitrogen-fixers and calcicole as they presented lower C/N than other species (p=0.0317, GLM, gaussian distribution, results not shown). High δ13C in leaves allows more time of stomatal closure causing higher water use efficiency (Goldstein et al. 1989; Scartazza et al. 1998). Therefore, our results show that nitrogen-fixing species have higher δ^13^C possibly as a consequence of stomatal closure responding to water limitation in soil followed by an intense CO2 depletion in N-enriched leaves pumping ^13^C into photosynthesis, a trait of plants that tolerate longer periods of water stress (Werner and Máguas 2010; Máguas et al. 2011). Thus, higher δ^13^C values suggest longer periods of biomass incorporation (Goldstein et al. 1989; Scartazza et al. 1998; Coletta et al. 2009) as well as different phenological developments (Werner and Máguas 2010) in tree species of Cerrado and show that δ^13^C variation is associated to N%, C% and C/N variation.

The relative disassociation of the SLA and stem elongation variation (driven by CEC and aluminium variation) from the N% and C% variation in leaves (associated to the variation soil inorganic nitrogen) appears in RLQ. However, stem elongation is also influenced by leaf N% and C/N according to GLMs indicating that all functional traits may be more or less associated in their variations responding to the main soil variables. However, the relative disassociation unveiled by the fourth-corner analysis can be a result of high water use efficiency of leguminous nitrogen-fixing species as they are N-enriched plants even in the Cerrado acidic soils with high aluminium saturation.

## Supporting information

Abundance data

## Acknowledgements

Authors thank the Botany Graduate Program and the Ecology Graduate Program of Universidade Federal de Viçosa. The authors also thank the Paraopeba Reserve staff for providing help and facilities.

