## Appendix S1 for "Responses of tree species traits to soil variation in the cerrado"

### Electronic Supplementary Material

### Appendix S1. Functional trait values of 34 most abundant species of the Cerrado at Paraopeba Reserve, Minas Gerais. (LEG = functional group of nitrogen-fixing leguminous species, OTR, other Cerrado species); SLA = specific leaf area (mm².mg^-1^); Hmax/Dmax = ratio of maximum height to maximum diameter; C/N = leaf carbon:nitrogen ratio; Bark = bark thickness (mm), Wood = wood density (mg.mm^-3^), NiTotal = abundance of species for the whole community; %abund = relative abundance of species). Wood density was gathered from an online database (Chav*e et al*., 2009), and so is the only trait not averaged. Plot specific calculation: for models’ calculation, each species had their individuals counted in a given plot and functional trait values of each species were multiplied by the number of individuals of each species in that given plot; then, the values found for each species were summed up for each functional group for that given plot

| Species | Family | Functional group | SLA | Hmax/Dmax | C/N | Bark | Wood | NiTotal | %abund |
| --- | --- | --- | --- | --- | --- | --- | --- | --- | --- |
| *Alibertia edulis* | Rubiaceae | OTR | 7.41 | 0.43 | 32.54 | 0.86 | 0.76 | 1981 | 0.1350 |
| *Astronium fraxinifolium* | Anacardiaceae | OTR | 5.25 | 0.97 | 22.81 | 1.68 | 0.82 | 159 | 0.0108 |
| *Bowdichia virgilioides* | Leguminosae | LEG | 3.01 | 0.54 | 20.08 | 3.30 | 0.79 | 163 | 0.0111 |
| *Byrsonima crassifolia* | Malpighiaceae | OTR | 7.21 | 0.27 | 34.84 | 1.94 | 0.58 | 370 | 0.0252 |
| *Callisthene major* | Vochysiaceae | OTR | 5.41 | 0.47 | 28.37 | 1.44 | 0.73 | 195 | 0.0133 |
| *Copaifera langsdorffii** | Leguminosae | LEG | 4.91 | 0.48 | 22.30 | 1.44 | 0.60 | 128 | 0.0087 |
| *Curatella americana* | Dilleniaceae | OTR | 6.96 | 0.39 | 32.64 | 1.91 | 0.65 | 139 | 0.0095 |
| *Dilodendron bipinnatum* | Sapindaceae | OTR | 8.38 | 0.79 | 20.76 | 1.94 | 0.69 | 101 | 0.0069 |
| *Erythroxyllum daphnites* | Erythroxyllaceae | OTR | 7.80 | 0.35 | 23.29 | 1.21 | 0.71 | 449 | 0.0306 |
| *Eugenia dysenterica* | Myrtaceae | OTR | 5.27 | 0.35 | 34.99 | 3.95 | 0.73 | 378 | 0.0258 |
| *Kielmeyera coriacea* | Clusiaceae | OTR | 3.52 | 0.47 | 40.83 | 4.52 | 0.29 | 269 | 0.0183 |
| *Leptolobium dasycarpum* | Leguminosae | LEG | 4.96 | 0.58 | 21.55 | 4.81 | 0.78 | 197 | 0.0134 |
| *Lithraea molleoides* | Anacardiaceae | OTR | 7.00 | 0.55 | 30.34 | 1.24 | 0.51 | 111 | 0.0076 |
| *Luehea divaricata* | Malvaceae | OTR | 9.05 | 0.76 | 28.03 | 2.78 | 0.56 | 461 | 0.0314 |
| *Machaerium opacum* | Leguminosae | LEG | 5.01 | 0.54 | 18.48 | 3.04 | 0.80 | 109 | 0.0074 |
| *Magonia pubescens* | Sapindaceae | OTR | 6.70 | 1.06 | 20.31 | 2.36 | 0.77 | 251 | 0.0171 |

***Appendix S1, cont.***

***Appendix S1, cont.***

| Species | Family | Functional group | SLA | Hmax/Dmax | C/N | Bark | Wood | NiTotal | %abund |
| --- | --- | --- | --- | --- | --- | --- | --- | --- | --- |
| *Miconia albicans* | Melastomataceae | OTR | 6.01 | 0.40 | 31.45 | 0.66 | 0.61 | 948 | 0.0646 |
| *Myracrodruon urundeuva* | Anacardiaceae | OTR | 8.13 | 0.50 | 23.99 | 1.80 | 1.00 | 207 | 0.0141 |
| *Myrcia tomentosa* | Myrtaceae | OTR | 6.36 | 0.84 | 23.88 | 1.26 | 0.80 | 203 | 0.0138 |
| *Myrsine umbellata* | Primulaceae | OTR | 6.11 | 0.59 | 28.57 | 1.55 | 0.49 | 372 | 0.0254 |
| *Pera glabrata* | Peraceae | OTR | 6.09 | 0.52 | 37.44 | 0.65 | 0.67 | 559 | 0.0381 |
| *Plathymenia reticulata* | Leguminosae | LEG | 7.72 | 0.60 | 20.40 | 2.32 | 0.49 | 67 | 0.0046 |
| *Platypodium elegans* | Leguminosae | LEG | 8.59 | 0.49 | 13.51 | 1.55 | 0.75 | 452 | 0.0308 |
| *Qualea grandiflora* | Vochysiaceae | OTR | 4.52 | 0.46 | 33.92 | 3.61 | 0.63 | 405 | 0.0276 |
| *Qualea parviflora* | Vochysiaceae | OTR | 5.83 | 0.33 | 30.86 | 4.12 | 0.63 | 372 | 0.0254 |
| *Roupala montana* | Proteaceae | OTR | 5.95 | 0.48 | 45.93 | 1.66 | 0.73 | 227 | 0.0155 |
| *Rudgea viburnoides* | Rubiaceae | OTR | 4.98 | 0.45 | 27.90 | 1.79 | 0.64 | 244 | 0.0166 |
| *Salvertia convallariodora* | Vochysiaceae | OTR | 3.58 | 0.20 | 33.32 | 6.03 | 0.65 | 37 | 0.0025 |
| *Styrax camporum* | Styracaceae | OTR | 5.97 | 0.36 | 29.63 | 1.49 | 0.34 | 241 | 0.0164 |
| *Tabebuia roseo-alba* | Bignoniaceae | OTR | 5.82 | 0.83 | 21.72 | 2.47 | 0.78 | 65 | 0.0044 |
| *Tapirira guianensis* | Anacardiaceae | OTR | 3.90 | 0.39 | 36.42 | 1.15 | 0.46 | 368 | 0.0251 |
| *Terminalia argentea* | Combretaceae | OTR | 5.88 | 0.54 | 32.96 | 1.47 | 0.81 | 232 | 0.0158 |
| *Vochysia tucanorum* | Vochysiaceae | OTR | 6.79 | 0.39 | 23.51 | 1.37 | 0.48 | 101 | 0.0069 |
| *Xylopia aromatica* | Annonaceae | OTR | 7.31 | 0.52 | 22.15 | 1.83 | 0.56 | 305 | 0.0208 |

* Species that is not confirmed as nitrogen-fixing sensu Sprent (2009).
